# Cytoplasmic zinc regulates IL-1β production by monocytes/macrophages via mTORC1-induced glycolysis in rheumatoid arthritis (RA)

**DOI:** 10.1101/2021.04.16.437150

**Authors:** Bonah Kim, Hee Young Kim, Bo Ruem Yoon, Jina Yeo, Kyung-Sang Yu, Hyeon Chang Kim, Jin Kyun Park, Seong Wook Kang, Won-Woo Lee

**Author notes:** Correspondence: Won-Woo Lee D.V.M., Ph.D., Professor, Department of Microbiology and Immunology, Department of Biomedical Sciences, Seoul National University College of Medicine, 103 Daehak-ro, Jongno-gu, Seoul 03080, South Korea., E-mail).

## Abstract

The essential micronutrient zinc plays regulatory roles in immune responses through its ability to affect signaling pathways. In activated monocytes/macrophages, signaling networks mediate metabolic reprogramming in order to meet the demands of participating in immune responses. Despite its known immunoregulatory roles, the effect of zinc on metabolic reprogramming in monocytes/macrophages remains unclear. Here, we demonstrate that cytoplasmic bioavailable zinc is essential for regulating IL-1β production in activated human monocytes/macrophages downstream of mTORC1-induced glycolysis. The cytoplasmic zinc level was influenced by extracellular zinc concentration through a zinc-specific importer, Zip8, which was markedly increased in monocytes of patients with rheumatoid arthritis (RA), a chronic inflammatory disease, and even in LPS-stimulated monocytes/macrophages of healthy individuals. Mechanically, phosphorylation of S6 kinase, a substrate of mTORC1, was significantly enhanced by zinc-mediated inhibition of PP2A, an S6 kinase phosphatase. As a result, IL-1β production was increased due to the activation of mTORC1-induced glycolysis. The expression of Zip8 and MT2A, a zinc-inducible gene, and the phosphorylation of S6 kinase by monocytes of RA patients was significantly enhanced compared with those of HCs and Zip8 levels positively correlated with RA clinical parameters, suggesting that Zip8-mediated zinc influx is related to inflammatory conditions. These results provide insight into the role of cytoplasmic bioavailable zinc in the metabolic reprogramming of human monocytes/macrophages which is an essential process for inflammatory responses.

**One Sentence Summary:** Cytoplasmic zinc regulates IL-1β production in monocytes/macrophages downstream of mTORC1-S6K-induced glycolysis *via* zinc-mediated inhibition of PP2A.

## Introduction

Zinc is an essential trace element that plays pivotal roles in multiple cellular functions(1, 2). It is well known for its conventional role as a cofactor that modulates structural or regulatory functions of thousands of proteins(3, 4). More recently, it has been suggested that zinc also functions as an intracellular signaling molecule, facilitating the transduction of a variety of signaling cascades in response to extracellular stimuli(5–8). Zinc deficiency is associated with various clinical problems including growth retardation, immune system dysfunction, and neurological disorders(9–11). Intracellular zinc homeostasis is tightly regulated by two families of proteins, the solute-linked carrier 39 (SLC39A, or Zip) family of zinc importers and solute-linked carrier 30 family (SLC30A, or ZnT) of zinc exporters. A total of 14 Zips and 10 ZnTs coordinately mediate flux of zinc ions across membranes in a cell- or tissue-specific manner(12).

Accumulating evidence demonstrates that zinc has an influence on the growth, development, and integrity of the immune system(1, 13). It has been demonstrated that abnormal zinc homeostasis caused by zinc deficiency impairs overall innate and adaptive immune functions. Zinc deficiency in the innate immune system is characterized by reduced PMN chemotaxis and phagocytosis of macrophages, with a resultant decrease in production of pathogen-neutralizing reactive oxygen species (ROS)(13). In addition, the production of pro-inflammatory cytokines by monocytes is markedly impaired by zinc deficiency(8, 14–16). In the adaptive system, zinc deficiency also detrimentally affects the development and function of T and B cells, which causes T-cell lymphopenia, imbalance among the different helper T-cell subsets, and reduced antibody production(17–19). Clinically, zinc deficiency resulting from malnutrition and dysregulated homeostasis increases susceptibility to viral and bacterial infections(20). Zinc supplementation has been reported to be beneficial for restoring immune function during various infectious diseases, including bacterial infections and malaria. However, mechanisms underlying zinc-mediated immune regulation and the resulting immunological consequences have not been well defined.

A number of studies have shown that activation-mediated zinc influx is spatiotemporally modulated in a variety of immune cells(6, 19, 21). In the cytoplasm bioavailable zinc ions participate in regulation of signaling molecule activity, which consequently shapes immune responses(1, 5). Mechanically, cytoplasmic zinc is known to regulate signal transduction through inhibition of phosphatases, including protein tyrosine phosphatases (PTPs) and serine/threonine phosphatases (PSPs), rather than by affecting kinase activity(22, 23). Given that phosphatases are generally involved in negative feedback of signaling activity, this aligns with the fact that cytoplasmic zinc ions elicit prolonged immune cell signaling and enhanced immune responses(6, 17, 18).

Inflammation is an indispensable process required to protect the host against pathogen invasion and tissue damage. Mononuclear phagocytes, especially monocytes and macrophages, play crucial roles in the initiation, regulation, and resolution of inflammatory responses through phagocytosis, cytokine production, generation of ROS, and activation of adaptive immunity(24, 25). Because immune responses are energy-demanding biosynthetic processes, stimulated monocytes and macrophages require intricate metabolic reprogramming to fulfill their metabolic requirements during immune responses. Among these, activation of glycolysis is a critical pathway in cellular metabolism that provides intermediates for energy generation(24). Recent studies have suggested that in response to the TLR4 agonist LPS there is increased glycolytic activity, which leads to increased production of inflammatory cytokines such as IL-1β. In macrophages the inhibition of glycolysis with 2-deoxy-D-glucose (2-DG) treatment markedly suppresses pro-IL-1β production and active IL-1β secretion(26). Several studies have suggested that the mammalian target of rapamycin complex 1 (mTORC1), a serine-threonine protein kinase, is both involved in aerobic glycolysis and regulates immune responses by monocytes/macrophages(24, 27, 28). Although zinc is a crucial factor involved in induction of IL-1β by monocytes/macrophages, little is known about its role in regulating metabolic reprogramming, especially during mTORC1-mediated glycolysis in human monocytes and macrophages.

Here, we examine the hypothesis that zinc functions as a regulator of IL-1β production in activated human monocytes/macrophages via mTORC1-induced glycolysis. Our data provide new insight into how cytoplasmic bioavailable zinc is involved in metabolic reprogramming of human monocytes/macrophages which is an essential process for inflammatory responses and might be is related to pathogenesis of inflammatory diseases such as RA.

## Results

### Enhanced expression of Zip8 in monocytes of patients with RA and activated monocytes/macrophages of HCs

Monocytes and macrophages are activated in RA patients(29–31), and thus, their unique regulation of zinc homeostasis is possibly involved in order to meet zinc demand. The re-analysis on our previous microarray data (E-MTAB-6187) showed that among 10 zinc-specific importer (ZnT) and 14 zinc-specific importer (Zip) proteins, expression of Zip8 was greatly upregulated in monocytes of RA patients when compared with those of healthy controls (Fig. 1A). This finding was confirmed in peripheral CD14^+^ monocytes purified from HCs and RA patients in a secondary cohort (Fig. 1B). Furthermore, mRNA expression of Zip8 was further enhanced in monocytes derived in synovial fluid, which is the site of inflammation in RA patients (Suppl. Fig. 1). Given activation-mediated changes in many zinc transporters by T cells(32), we first assessed the expression of Zip genes in monocytes and monocytes-derived macrophages (hereafter, macrophages) upon LPS stimulation. As seen in Figure 1C and D, resting monocytes and macrophages had relatively higher expression of Zip1 and Zip8 compared to other Zips. In addition, LPS stimulation led to a dramatic increase of Zip8 in both monocytes and macrophages. This suggests that Zip8 plays important roles in the regulation of zinc influx in monocytes and macrophages.

**Figure 1.**
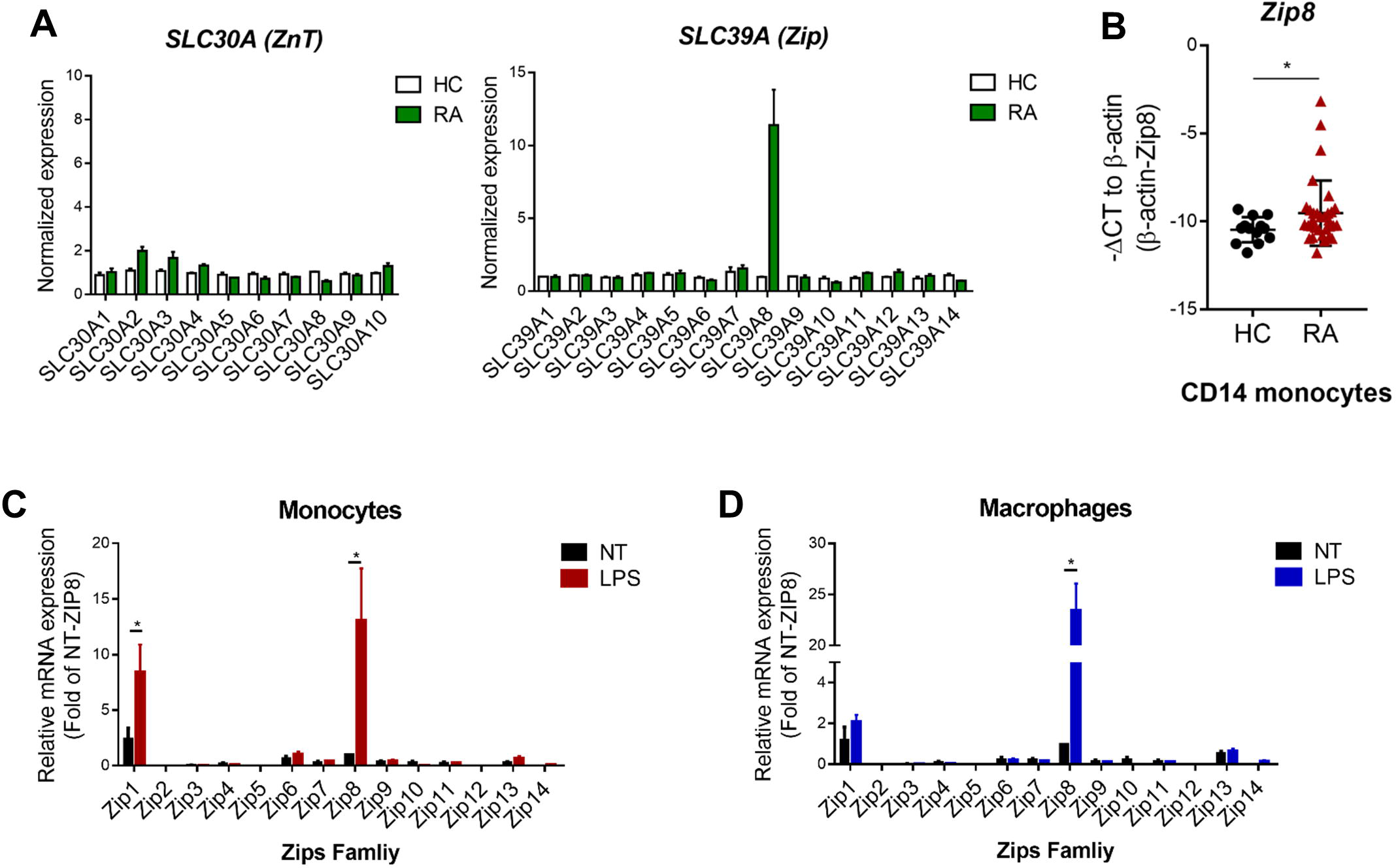
Enhanced expression of zinc transporter Zip8 in monocytes of patients with RA and activated monocytes/macrophages of HCs. **(A)** Microarray analysis on 10 SLC30A (ZnT) and 14 SLC39A (Zip) transporters expressed by peripheral monocytes. CD14^+^ monocytes were purified from peripheral blood mononuclear cells of RA patients (*n* = 3) and healthy controls (HCs) (*n* = 2). **(B)** Quantitative PCR analysis of Zip8 gene expression by peripheral monocytes derived from HC (*n* = 13) and RA patients (*n* =32). Expression was normalized to β-actin, and the comparative Ct method was used for the quantification of gene expression. (**C-D**) mRNA levels of 14 Zip family genes were quantified by real-time RT-PCR in monocytes (C) (n=7) and macrophages (D)(n=5) stimulated with or without LPS for 24 hr. Relative expression of mRNA of ZIP family genes was normalized to Zip8 without LPS. Bar graphs and scatter plots show the mean ± SEM. * = *p* < 0.05 by two-tailed paired *t*-test.

### Zinc influx is dependent on extracellular zinc levels and occurs via Zip8 transporters

To examine whether the increase in intracellular zinc in monocytes/macrophages is dependent on extracellular zinc, FluoZin-3, a zinc specific-fluorescent probe, was used to monitor cytoplasmic, bioavailable zinc ions in real-time. Although the pattern of cytoplasmic zinc increase differs between monocytes and macrophages, influx of zinc ions was found to occur immediately after treatment with FluoZin-3 buffer supplemented with zinc. Moreover, the intracellular zinc level was found to be dependent on the extracellular zinc concentration (Fig. 2A and B). This increase was sustained until 50 min after zinc treatment. Metallothionein (MT) is a metal-binding protein that plays coordinated roles in the distribution, transport, and maintenance of intracellular zinc(33). Further, the induction of MT expression is dependent on the increase of intracellular zinc. Our data show that the expression of MT2A, a major isoform of MT family proteins, was markedly induced in proportion to zinc influx in monocytes and macrophages. LPS stimulation further upregulated MT2A mRNA expression, suggesting that increased zinc influx occurs following LPS stimulation (Fig. 2C and D). Since the zinc transporter Zip8 is highly expressed in resting and activated human monocytes and macrophages (Fig. 1C and D), we tested whether Zip8 contributes to an increased zinc influx by a Zip8 siRNA (siZip8) knockdown experiment. Macrophages were transfected with Zip8-targeted or scrambled siRNA followed by stimulation with LPS for 18 h. As evidenced by qPCR and immunoblotting, expression of Zip8 in macrophages was efficiently silenced (over 80% reduced; Suppl. Fig. 2), and this led to a significant reduction of both zinc influx and MT2A mRNA induction (Fig. 2E and F). Lastly, we found that the cytoplasmic zinc level in *ex vivo* monocytes of healthy donors exhibits a significant positive correlation with the plasma level of zinc ions (Fig. 2G; *p* = 0.026), supporting our findings (Fig. 2A and B). These data demonstrate that zinc is fluxed into monocytes and macrophages in part via Zip8 and that this influx is dependent on extracellular zinc levels.

**Figure 2.**
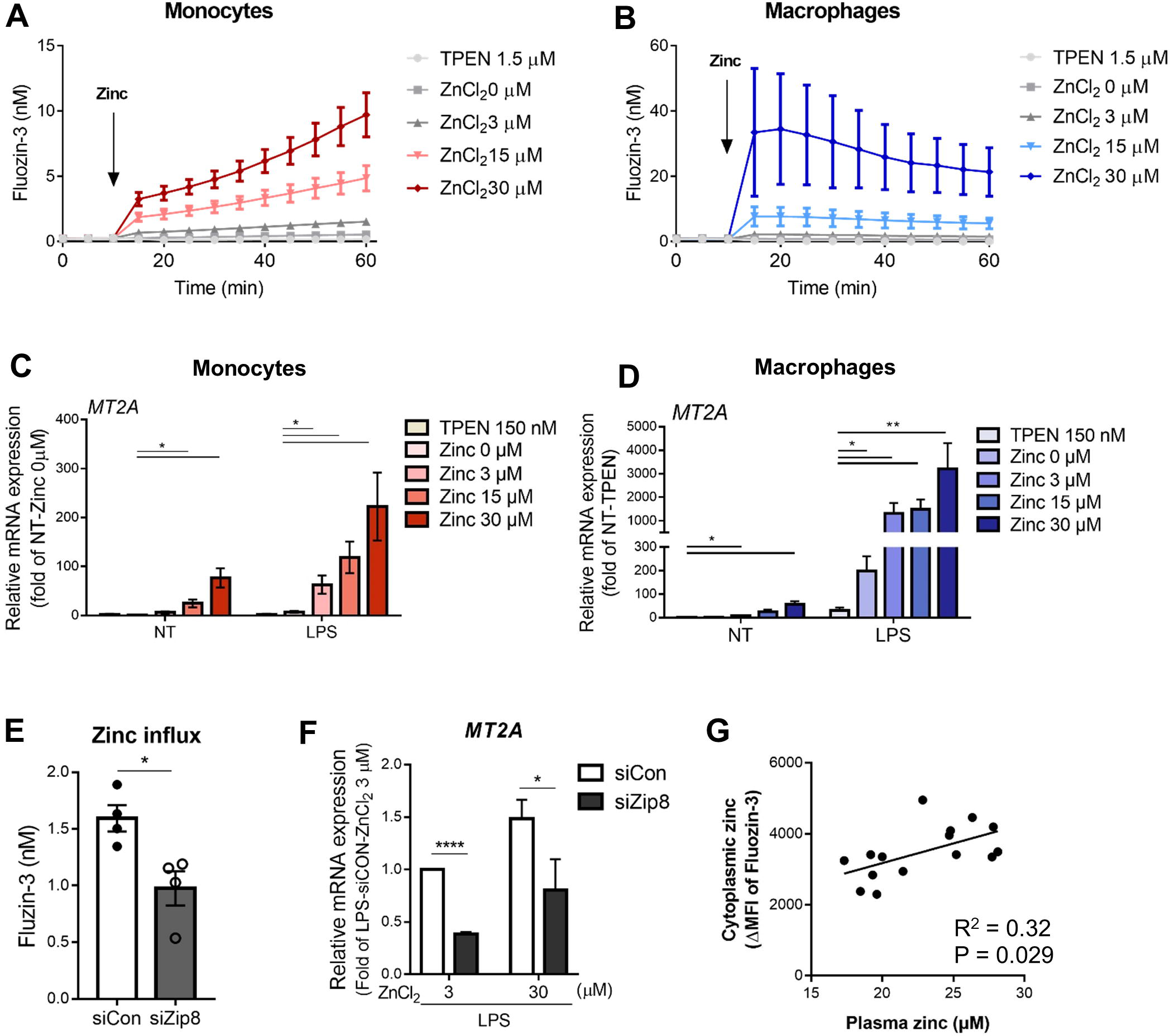
Zinc influx is dependent on extracellular zinc levels and occurs via Zip8 transporters. **(A-B)** Intracellular zinc was measured in human monocytes (A) and macrophages (B) loaded with FluoZin-3. After recording the baseline fluorescence for 10 min, different concentrations of ZnCl_2_ were added into the cells (arrow) and the FluoZin-3 signal was recorded for an additional 50 min. The concentration of zinc ions was calculated as described in Material and Methods. A representative experiment (mean of triplicates ± SD) of three independent experiments is shown. **(C-D)** MT2A mRNA level was quantified by qRT-PCR in monocytes (C) (n=5) and macrophages (D) (n=4) stimulated with LPS. **(E-F)** Macrophages were transfected with Zip8-targeted or control siRNA and then incubated with ZnCl_2_ for 2 hr, followed by stimulation with LPS (10 ng/ml) for 24 hr. (E) Influx of zinc ions was monitored by FluoZin-3. (F) Zip8 mRNA was quantified by qRT-PCR. **(G)** Correlation between plasma zinc (μM) and cytoplasmic bioavailable zinc level (AU) in monocytes of healthy donors (n = 15). *p* value was obtained using the Pearson correlation analysis. Bar graphs show the mean ± SEM. * = *p* < 0.05, ** = *p* < 0.01, and **** = *p* < 0.0001 by two-tailed paired *t*-test.

### Increased extracellular zinc boosts production of IL-1β in human monocytes/macrophages

To explore the role of zinc in the production of effector cytokines by monocytes and macrophages, we tested whether increased extracellular zinc influences the production of IL-1β following LPS stimulation of these cells. Freshly isolated monocytes and monocyte-derived macrophages were pretreated for 2 hr with TPEN (*N*,*N*,*N′*,*N′*-tetrakis(2-pyridinylmethyl)-1,2-ethanediamine), a membrane-permeant zinc-specific chelator, or culture media supplemented with different concentrations of ZnCl_2_, followed by stimulation with LPS for monocytes or LPS and ATP or macrophages. The viability of monocytes was not influenced by these culture conditions (Suppl. Fig. 3A). The secretion of IL-1β in the supernatant of LPS-stimulated monocytes and macrophages was potently increased in a zinc concentration-dependent manner (Fig. 3A and B). The increase in intracellular pro-IL-1β in LPS-stimulated monocytes and macrophages also occurred in a zinc concentration-dependent manner (Fig. 3C and 4D). Moreover, this effect was more apparent in the culture supernatant, as it showed that the proteolytic processing and secretion of caspase-1 and IL-1β resulted in increased levels of these cytokines in a zinc concentration-dependent manner. In addition to IL-1β, the production of TNF-α and IL-6 was also dependent on zinc concentration (Suppl. Fig. 3B) as previously reported(8). These data show that zinc is important for production of IL-1β and increased extracellular zinc promotes production of IL-1β in human monocytes/macrophages.

**Figure 3.**
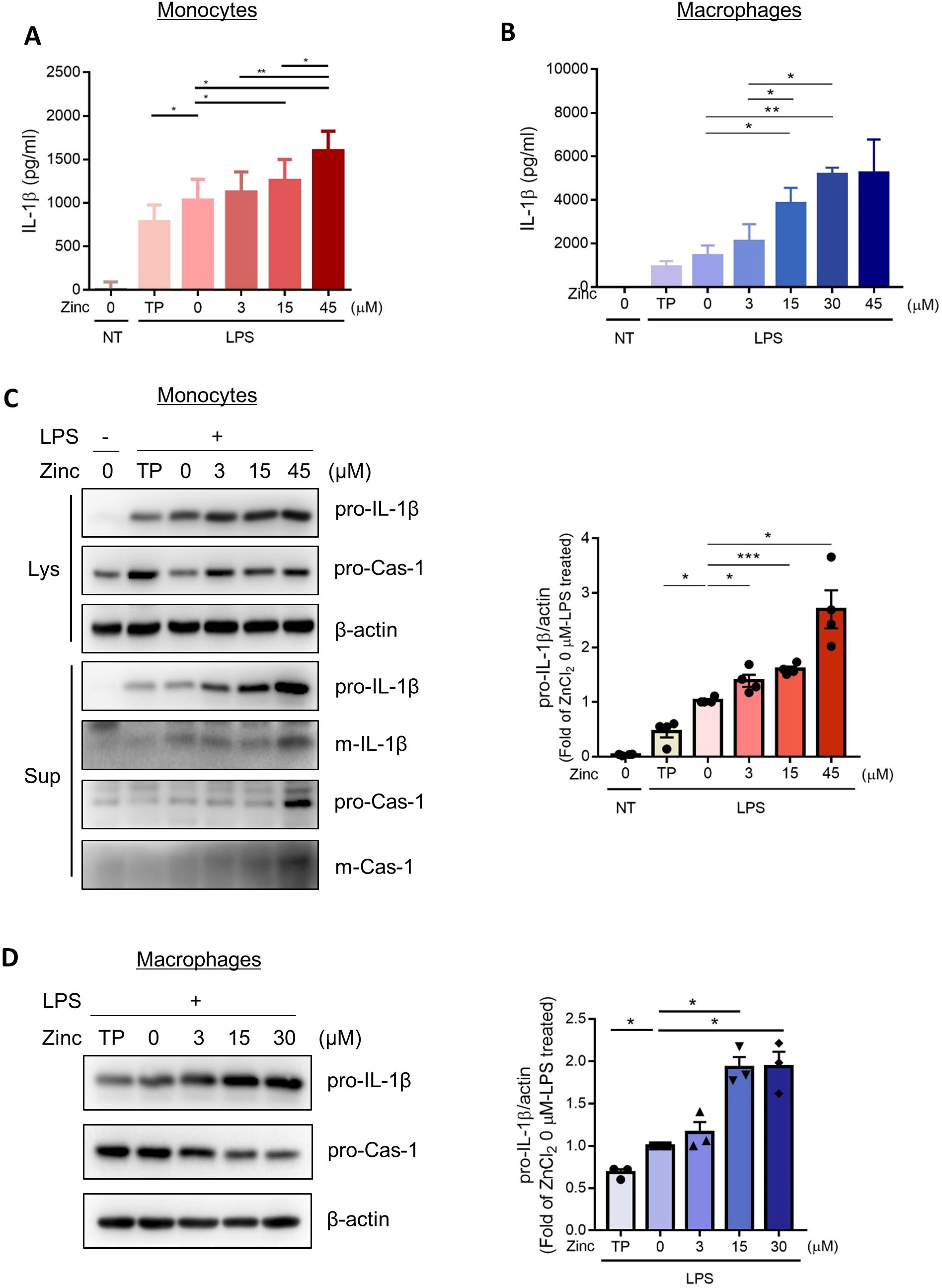
Increased extracellular zinc boosts production of IL-1β in human monocytes/macrophages. **(A-B)** Freshly purified monocytes from healthy donors (A) and monocyte-derived macrophages (B) were incubated with the zinc chelator TPEN (150 nM) or the indicated concentration of ZnCl_2_ for 2 hr, followed by stimulation with LPS for 24 hr. Macrophages were given additional stimulation with ATP for the last 6 h. The amount of IL-1β in the culture supernatants of monocytes (A; n=7) and macrophages (B; n=4) was quantified by ELISA. (**C-D**) Cell extracts (Lysate) and supernatants (Sup) of monocytes in (C) and cell extracts (Lysate) of macrophages in (D) were prepared for immunoblotting for IL-1β and caspase-1 proteins. Band intensity in immunoblots was quantified by densitometry. β-actin was used as a normalization control (n=3). Bar graphs show the mean ± SEM. * = *p* < 0.05, ** = *p* < 0.01, and *** = *p* < 0.005 by two-tailed paired *t*-test.

**Figure 4.**
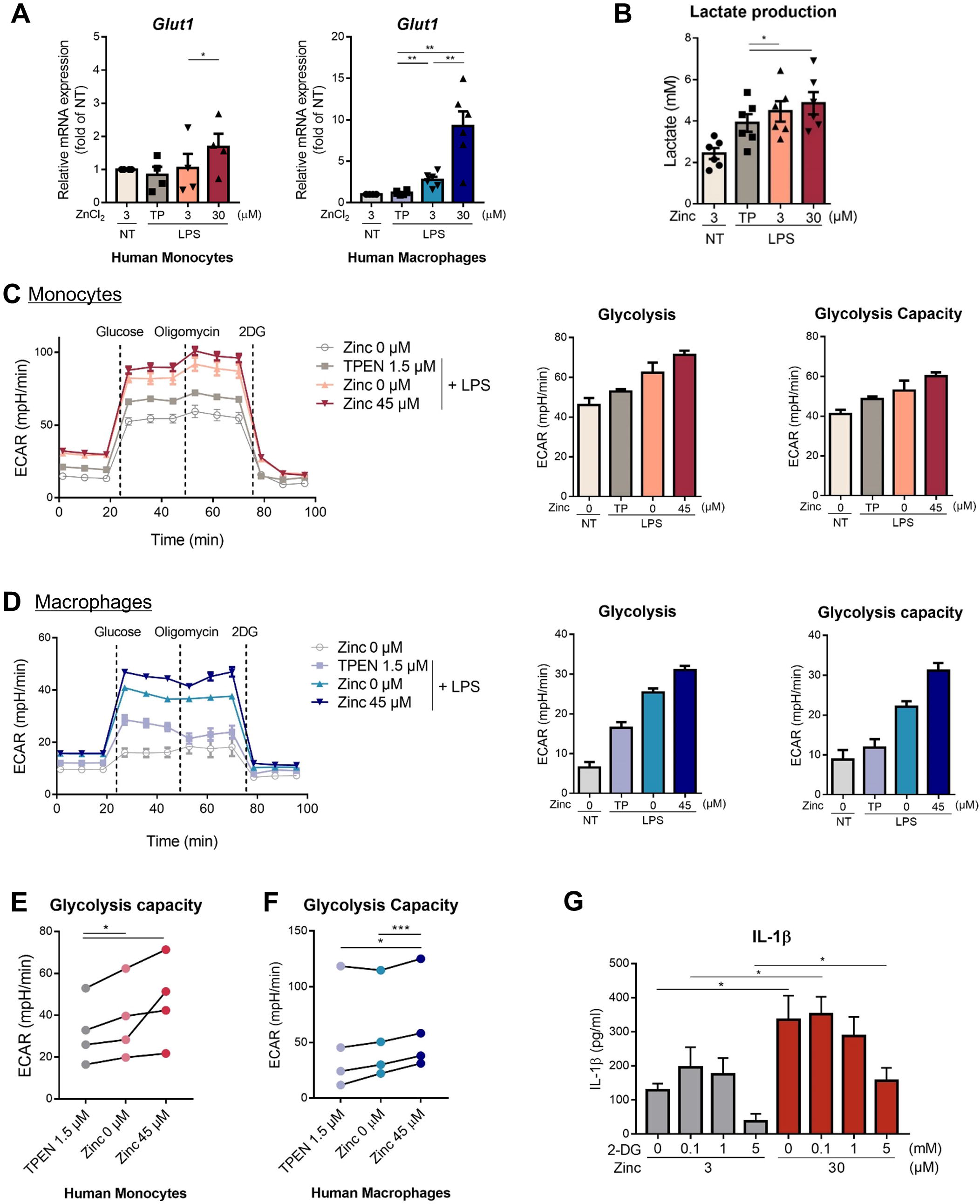
Increased intracellular zinc is associated with upregulation of glycolytic metabolism. **(A)** Monocytes and macrophages were treated with TPEN 150 nM or ZnCl_2_ for 2 hr, followed by stimulation with LPS for 24 hr. The mRNA expression of Glut1 was quantified by real-time RT-PCR. Relative expression of Glut1 was normalized to monocytes without LPS treatment. **(B)** The amount of lactate in the culture supernatant of monocytes was measured using the lactate colorimetric Assay Kit. **(C-D)** ECAR (extracellular acidification rate) was measured in monocytes (C) and macrophages (D) pre-incubated with TPEN (1.5 μM) or ZnCl_2_ (0 or 45 μM) for 2 hr and stimulated with LPS for 24 hr. ECAR levels were measured following sequential treatment with glucose, oligomycin, and 2-DG. (**E-F**) Cellular glycolysis capacity in LPS-stimulated monocytes (E) and macrophages (F) under the indicated zinc concentrations. Monocytes and macrophages from four different donors were independently tested. **(G)** Monocytes were pre-incubated with the indicated concentration of 2-DG and ZnCl_2,_ followed by stimulation with LPS for 4 hr. The amount of IL-1β in culture supernatant was quantified by ELISA (n=6). Bar graphs show the mean ± SEM. * = *p* < 0.05, ** = *p* < 0.01, and *** = *p* < 0.005 by two-tailed paired *t*-test.

### Increased intracellular zinc is associated with upregulation of glycolytic metabolism

In innate immune cells glycolytic metabolism is closely linked to the production of immune-critical cytokines such as IL-1β(27, 34, 35). Enhanced glucose influx, which is facilitated by upregulated expression of glucose transporter 1 (GLUT1), is intimately associated with their glycolytic stimulation(36). We found that Glut1 expression is significantly increased in LPS-stimulated monocytes and macrophages under higher zinc concentrations (Fig. 4A). Furthermore, lactate, a glycolytic end-product, was also increased in human monocytes in proportion to the cytoplasmic bioavailable zinc level compared to that of the unstimulated control (Fig. 4B). To further investigate the effect of zinc on glycolytic metabolism in activated monocytes and macrophages, the extracellular acidification rate (ECAR), a parameter of glycolytic metabolism, was measured in monocytes or macrophages activated by LPS in the presence of different concentrations of zinc or TPEN. As expected, LPS stimulation induced glycolytic metabolism in monocytes and macrophages (Fig. 4C and D). The ECAR in monocytes and macrophages was markedly increased by higher levels of zinc compared to 3 μM zinc or TPEN treated groups (Fig. 4C-F). To functionally explore the effect of zinc on glucose metabolism during glycolysis, monocytes were treated with 2-DG, a glucose analog, to inhibit glucose metabolism. Treatment with 30 μM zinc only partially antagonized the inhibitory effect of 2-DG on IL-1β production, showing that IL-1β production of monocytes treated with 30 μM zinc and 5 mM of 2-DG was comparable that of monocytes treated with 3 μM zinc without 2-DG (Fig. 4G). This data suggests that enhanced glycolytic metabolism in human monocytes/macrophages might be affected by increased intracellular zinc.

### Intracellular zinc is important for activation of the mTORC1-S6K signaling pathway

Zinc plays a fundamental role in controlling monocyte/macrophage functions and intracellular zinc is essential for activation-induced signal transduction in monocytes(1, 8, 37–39). As reported(8, 40), phosphorylation of Erk1/2 was found to markedly increase in a zinc concentration-dependent manner in activated human primary monocytes, whereas zinc had no obvious effect on NF-κB or p38 activity (Suppl. Fig. 4A). Similar to Erk1/2, the phosphorylation of mTOR activity was upregulated with treatment under conditions of increased zinc, suggesting zinc affects more than one major signaling pathway in activated monocytes (Suppl. Fig. 4A and Fig. 5A). Others and we reported that mTORC1-S6K activation is responsible for aerobic glycolysis that plays a pivotal role in the effector function of monocytes/macrophages (27, 28, 41). Thus, we sought to explore the signaling role of zinc in the mTOR pathway. To this end, phosphorylation of upstream and downstream molecules in the mTOR pathway, such as Akt, S6K and 4E-BP1, were analyzed in zinc-treated monocytes. Chelation of zinc with TPEN caused remarkably diminished Akt/mTOR activation in monocytes. Moreover, we found that cytoplasmic zinc is indispensable for phosphorylation of S6K, a substrate of mTORC1 (Fig. 5B). To further elucidate whether zinc-dependent enhancement of S6K activity is simply a downstream consequence of increased Akt activity, monocytes were treated with MK-2206, an Akt-selective inhibitor, and incubated with culture media supplemented with 3 or 30 μM of zinc. As seen in Figure 5C, phosphorylation of Akt was dramatically inhibited in the MK2206-treated monocytes, irrespective of zinc concentration, whereas the inhibitory effect of S6K activation by MK2206 was limited in monocytes treated with 30 μM of zinc. Figure 5D shows that enhancement of Akt activation by 30 μM zinc treatment is completely inhibited by 10 nM MK2206, while S6K activation is not significantly changed under these conditions, suggesting involvement of other mechanisms in zinc-dependent enhancement of S6K activity. A similar finding was made in LPS-activated monocytes (Suppl. Fig. 4B). Of interest, zinc-mediated phosphorylation of S6K was observed in monocytes and macrophages even without LPS stimulation (Fig. 5E). Since phosphorylation of the ribosomal S6 protein (S6) is directly linked to the activity of the mTORC1-S6K(42), we analyzed the relationship between phosphorylation of S6 and the level of cytoplasmic bioavailable zinc in *ex vivo* monocytes from healthy donors and found a significant correlation (Fig. 5F; *p* = 0.0036). As seen in Figure 5G and H, LPS stimulation markedly enhanced mTORC1-S6K activation in a time-dependent manner until 1 hr post-activation. Further, this activity was maintained until 6 hr post-activation in LPS-stimulated macrophages (Suppl. Fig. 4C). Since glucose metabolism is critical for a shift towards glycolytic reprogramming, 2-DG abated phosphorylation of S6K in a dose-dependent manner (Fig. 5I). However, in agreement with our finding in Figure 4G, treatment with 30 μM zinc partially antagonized the inhibitory effect of 2-DG on S6K activity (Fig. 5J), showing that phosphorylation of S6K in monocytes treated with 30 μM zinc and 5 mM of 2-DG was comparable that of monocytes treated with 3 μM zinc without 2-DG. Our data demonstrate that intracellular zinc is critical for mTORC1-S6K activation in human monocytes and macrophages and zinc-mediated mTORC1 activity provokes enhanced glycolytic metabolism.

**Figure 5.**
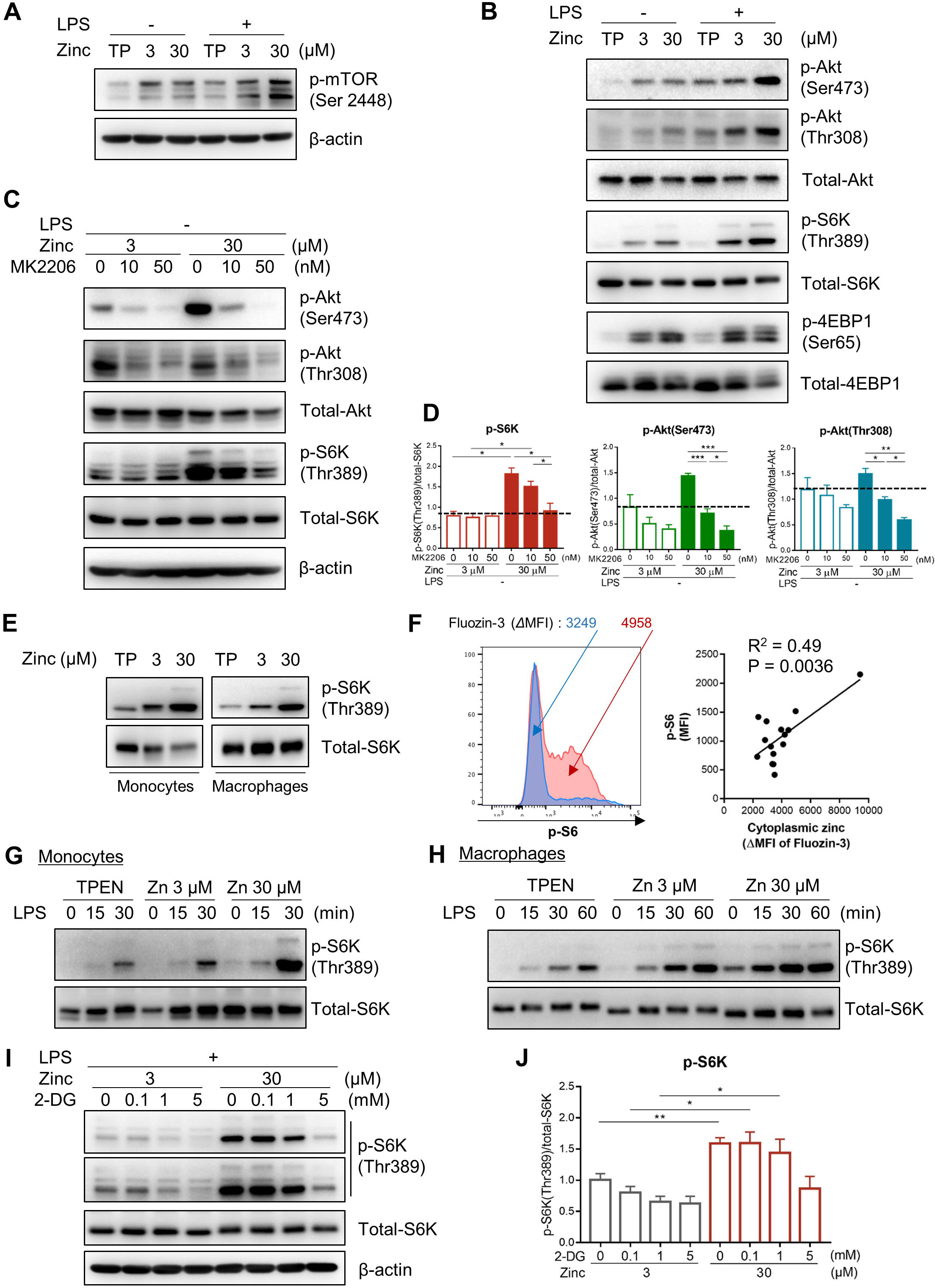
Intracellular zinc is important for activation of the mTORC1-S6K signaling pathway. **(A-B)** Immunoblot analysis on Akt-mTOR-S6K pathway in monocytes were treated with TPEN or ZnCl_2_ for 2 hr and stimulated with or without LPS for 15 min (n=3). **(C)** Monocytes were pre-treated with Akt inhibitor, MK2206, followed by incubation with ZnCl_2_ (n=4). **(D)** The phosphorylation of Akt and S6K in monocytes (C) was plotted with band intensities. The phosphorylation level was normalized to the expression of total form protein. **(E)** Cells were treated with TPEN or ZnCl_2_ for 30 or 15 min, respectively. **(F)** A representative histogram plot of phosphorylated S6 in *ex vivo* CD14^+^ monocytes from healthy donors (left). Correlation of p-p70-S6K with cytoplasmic zinc level (AU) in monocytes of healthy donors (n = 15) (right). *p* value was obtained using the Pearson correlation analysis. **(G-H)** Cells were pre-incubated with TPEN or ZnCl_2_ for 2 hr, followed by LPS stimulation for the indicated time. (**I)** Monocytes were pre-incubated with 2-DG and ZnCl_2,_ followed by LPS stimulation. Data is representative of three independent experiments. **(J)** Band intensity of p-p70-S6K in immunoblots (I) was quantified. Bar graphs show the mean ± SEM. * = *p* < 0.05, ** = *p* < 0.01, and *** = *p* < 0.005 by two-tailed paired *t*-test.

### The effects of zinc on mTORC1 activity are mediated by PP2A in human monocytes/macrophages

Our data thus far suggest that the zinc-mediated activation of the mTORC1-S6K pathway contributes to increased production of IL-1β via enhanced glycolytic metabolism. Next, we sought to investigate the molecular mechanisms underlying the regulation of phosphorylation of S6K by zinc. Zinc ions are known to inactivate several types of protein phosphatases(6, 18, 22). Therefore, we hypothesized that cytoplasmic zinc ions influence the activity of protein phosphatases regulating mTORC1-S6K activity in human monocytes and macrophages. Recent studies have shown that S6K activity is negatively regulated by serine/threonine phosphatases, such as protein phosphatase 2A (PP2A) and pleckstrin homology domain leucine-rich repeat protein phosphatase (PHLPP), via dephosphorylation(43, 44). However, little is known about the effect of zinc on these protein phosphatases in immune cells. PP2A protein, including PP2A-B and C subunits, is abundantly expressed in *ex vivo* monocytes with no obvious change in expression observed following LPS stimulation (Fig. 6A). Treatment with LB100, a small molecule inhibitor of PP2A, was found to intensify LPS-induced phosphorylation of mTOR pathway-related molecules, including Akt, mTORC1 and S6K, in human monocytes (Fig. 6B). Furthermore, IL-1β production was significantly increased by inhibition of PP2A activity with LB100 (Fig. 6C). Similar results were also obtained in LPS-stimulated monocytes treated with Okadaic acid, a general inhibitor of serine/threonine phosphatases PP1 and PP2A (Fig. 6C and Suppl. Fig. 5A). Since PP2A is known to interact with its substrates, we examined its physical associations in cells. The interaction of PP2A subunits with S6K was detected in transiently transfected 293T cells (Fig. 6D) by immunoprecipitation. More importantly, the interaction between endogenous PP2A and S6K was confirmed in resting and LPS-stimulated human macrophages (Fig. 6E). Given that phosphorylation of Tyr307 of the PP2A-C subunit decreases its phosphatase activity(45, 46), we next asked whether zinc-enhanced phosphorylation of S6K is attributable to inhibition of phosphatase activity through this mechanism. To this end, phosphorylation of PP2A was examined in monocytes incubated with different zinc concentrations. Treatment with 30 μM of zinc caused significantly increased phosphorylation of the PP2A-C subunit compared to monocytes treated with TPEN or 3 μM of zinc (Fig. 6F). PP2A activity was also measured by *in vitro* serine/threonine protein phosphatase assay to investigate the direct effect of zinc on phosphatase activity. Lysates, which were prepared from primary monocytes, were treated with the indicated concentration with ZnCl_2_, followed by reaction with a chemically synthesized phosphopeptide substrate specific for PP2A, but not PP1. As seen in Figure 6G, at concentrations over 500 μM zinc significantly inhibited PP2A activity. The monocyte lysates treated with 1,000 μM of zinc had PP2A activity as low as those treated with LB100, the PP2A-specific inhibitor. It should be noted that the effective zinc concentration in the *in vitro* phosphatase assay was supraphysiological compared to its concentration in our cell culture system. Because various buffers used for the *in vitro* phosphatase assay may affect the bioavailable zinc concentration (possibly via chelation), we measured the actual zinc ion concentration using a conventional zinc assay kit. No zinc ion was detected in lysates from cells treated with less than 100 μM of zinc, whereas the zinc level was around 3 μM in the lysates of cells treated with 500 μM zinc, which exhibited an inhibitory effect on PP2A activity in cell culture, suggesting there was zinc-chelating activity of the buffers used in the phosphatase assay (Suppl. Fig. 5B). Lastly, we investigated whether PHLPP, another candidate phosphatase, is involved in zinc-mediated regulation of S6K activity and glycolysis. PHLPP was not found to interact with S6K in THP-1 derived macrophages (Suppl. Fig. 6A) nor was the PHLPP-specific inhibitor, NSC-45586, found to have an effect on the phosphorylation of S6K in LPS-stimulated monocytes (Suppl. Fig. 6B). Thus, there appears to be no regulatory role of PHLPP on S6K activity in human monocytes and macrophages. These data demonstrate that the regulation of mTORC1 activity by PP2A phosphatase is medicated by intracellular zinc in human monocytes and macrophages.

**Figure 6.**
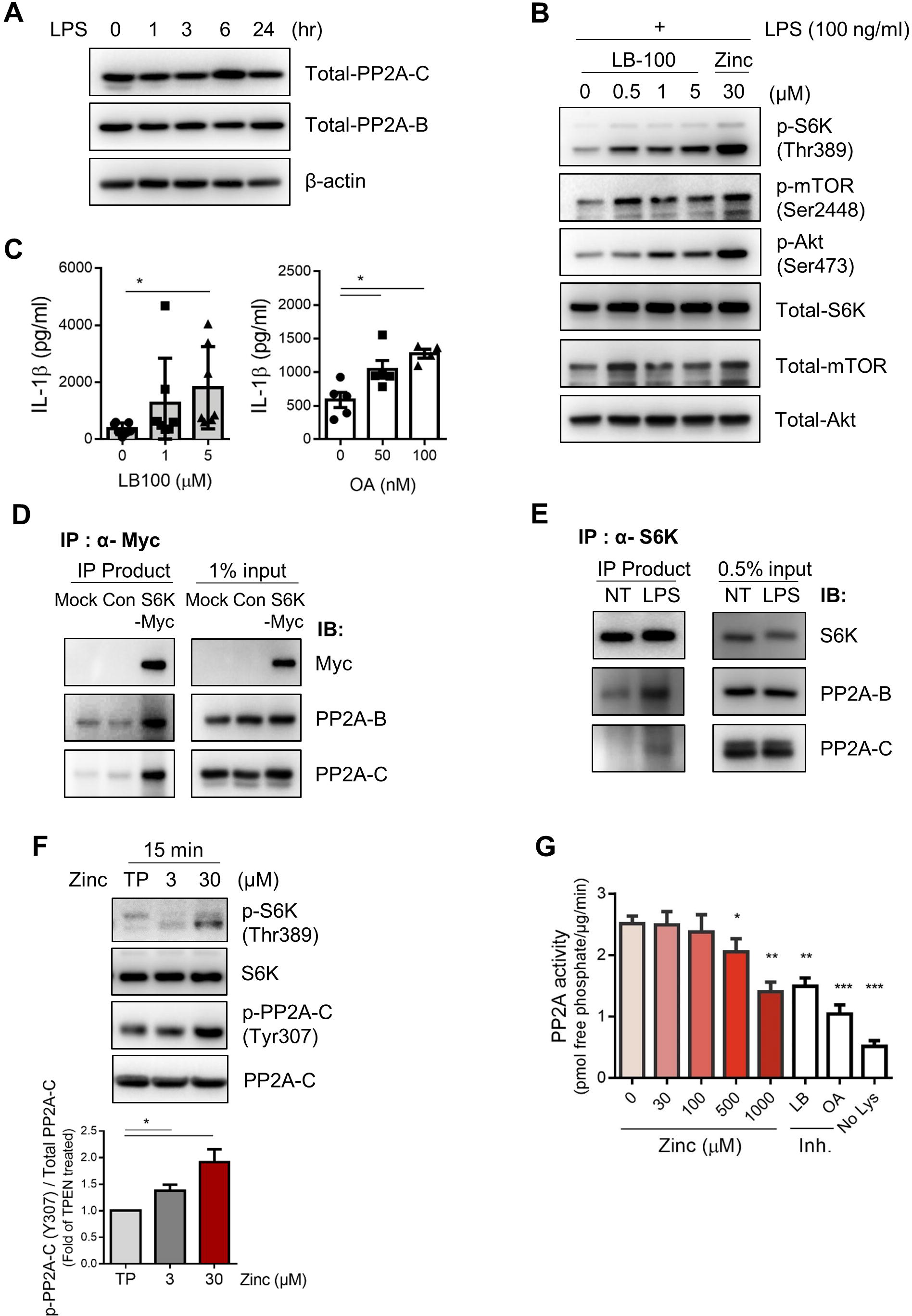
The effects of zinc on mTORC1 activity are mediated by PP2A in human monocytes/macrophages. **(A)** Monocyte cell lysates were prepared at 0, 1, 3, 6, 24 hr after stimulation with LPS (100 ng/ml). **(B)** Monocytes were pretreated with LB-100 or ZnCl_2_ for 2 hr, followed by stimulation with LPS for 30 min. **(C)** The amount of IL-1β was quantified by ELISA in the culture supernatants of LPS-stimulated monocytes in the presence of LB-100 or Okadaic acid. **(D-E)** 293T cells were transiently transfected with Myc-p70 S6K or control plasmid vector. Cell lysates of 293T and human macrophages were immunoprecipitated (IP) with antibodies to Myc or S6K and immunoblotted with the indicated antibodies. **(F)** Immunoblot analysis of phosphorylation of p70-S6K and PP2A-C was performed with monocytes, which were treated with TPEN (150 nM) and ZnCl_2_ (3 or 30 μM) for 15 min. **(G)** Lysates from freshly purified monocytes were incubated with various concentration of ZnCl_2_ (0, 30, 100, 500 or 1,000 μM), LB-100 (5 μM), or Okadaic acid (50 nM). PP2A activity was measured using a protein phosphatase activity assay kit as described in Material and Methods. Bar graphs show the mean ± SEM. * = *p* < 0.05, ** = *p* < 0.01, and *** = *p* < 0.005 by two-tailed paired *t*-test.

### Zinc-mediated metabolic reprogramming in monocytes is associated with clinical parameters of RA patients

Our data thus far suggest that Zip8-mediated influx of zinc contributes to increased production of IL-1β by activated mTORC1-induced glycolysis in monocytes and macrophages. Sustained hyper-inflammatory activity of glycolytic monocytes/macrophages is a key parameter of RA(24, 47), suggesting that metabolic reprogramming occurs in these cells. Therefore, we sought to examine whether zinc-mediated metabolic reprogramming is associated with clinical parameters and disease activity of RA patients. Zip8 expression by monocytes of RA patients had a significant positive correlation with disease activity score 28 based on erythrocyte sedimentation rate (DAS28-ESR) and C-reactive protein (DAS28-CRP), which represent enhanced inflammatory responses (Fig. 7A, *p* = 0.0072 and *p* = 0.0050, respectively and Suppl. Fig. 7A). Peripheral monocytes of RA patients had significantly higher mRNA level of MT2A, which is directly induced by cytoplasmic zinc, compared with HCs (Fig. 7B). The increase of MT2A was amplified in synovial monocytes of RA patients (Suppl. Fig. 7B). Furthermore, mRNA level of MT2A in monocytes of RA patients was significantly correlated with Zip8 and IL-1β expression (Fig. 7C), suggesting that zinc-mediated metabolic reprogramming may contribute to establishment of the inflammatory milieu in RA disease. Consistent with *in vitro* findings, *ex vivo* peripheral monocytes derived from RA patients had significantly increased phosphorylation of S6K and showed increased trend of intracellular pro-IL-1β (Fig. 7D and E). More importantly, PP2A activity of lysates, which were prepared from primary monocytes, was reduced in RA patients compared with HCs (Fig. 7F). This finding corroborates an increased activity of S6K in monocytes of RA patients (Fig. 7D). To further investigate the relevance between Zip8 expression and S6K-mediated IL-1β production, we compared the clinical parameters, levels of MT2A expression, phosphorylation of S6K, and pro-IL-1β in monocytes derived from peripheral monocytes of patients sets with low or high Zip8 expression. There was a significant increase in disease activity within the Zip8 high-expression group (Fig 7G). Moreover, RA monocytes with elevated Zip8 expression had significantly increased MT2A expression and phosphorylation of S6K, and showed increasing trend of intracellular pro-IL-1β (Fig. 7H-J). Together, these results suggest that the enhanced zinc influx by Zip8 in monocytes from RA patients plays a role in the regulation of inflammatory responses in RA.

**Figure 7.**
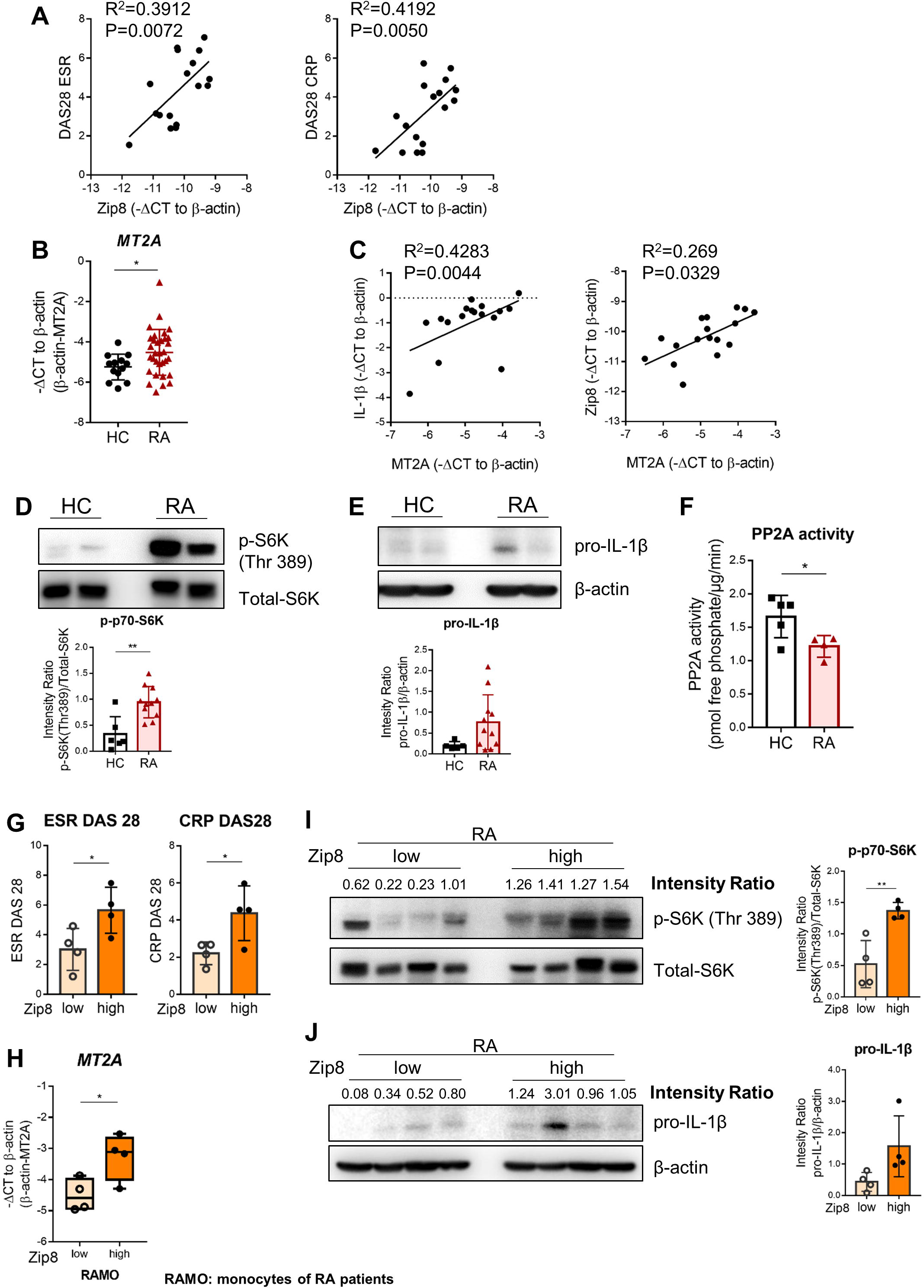
Zinc-mediated metabolic reprogramming in monocytes is associated with clinical parameters of RA patients. **(A)** Correlation of Zip8 gene expression in peripheral monocytes with RA clinical parameters (*n* = 17). **(B)** mRNA level of MT2A was quantified by real-time RT-PCR in peripheral monocytes of HCs (*n*=13) and RA patients (*n*=32). (**C**) Correlation of MT2A gene expression in peripheral monocytes with Zip8 or IL-1β gene expression in RA patients (*n* = 17). (**D-**Immunoblot analysis of phosphorylation of p70-S6K and pro-IL-1β was performed with monocytes of HCs (*n* = 6) and RA patients (*n* = 11). (**F**) PP2A activity was measured in lysates from freshly purified monocytes of HCs and RA patients as described in Figure 6G. (**G-H**) *Δ*Ct of Zip8 mRNA in the two groups (Mean ± SEM; −9.58±0.15 of low and −8.31±0.67 of high group). Scatter plots show RA clinical parameters and MT2A mRNA level in monocytes of RA patients having higher and lower zip8 mRNA expression. **(I-J)** Immunoblot analysis of phosphorylation of p70-S6K and pro-IL1β were performed with monocytes of RA patients having higher and lower Zip8 mRNA expression. *p* values were obtained using the Pearson correlation analysis (A and C). Bar graphs and scatter plots show the mean ± SD. * = *p* < 0.05 and ** = *p* < 0.01 by two-tailed unpaired *t*-test.

## DISCUSSION

Two different types of intracellular zinc signaling have been proposed during immune responses. The first type is mediated by a prompt increase of intracellular zinc upon extracellular stimulation as described in FcεR-stimulated mast cells and TCR-activated T cells(6, 7). In contrast, another type of zinc signaling is induced several hours after stimulation and depends on transcriptional changes in zinc transporter expression as depicted in LPS-stimulated dendritic cells (DCs) and TCR-activated T cells(18, 21, 48). In our study, zinc influx into human monocytes was found to be mediated within 10 min of LPS stimulation in a transcription-independent manner as previously reported (Fig. 2A and B)(8, 15). Meanwhile, the increase in cytoplasmic zinc is further exacerbated by LPS-mediated induction of Zip8 expression on the plasma membrane (Fig. 2C and and this leads to increased IL-1β production downstream of the mTORC1-S6K pathway (Fig. 3A, B, and 5B).

Intracellular zinc homeostasis is exquisitely governed by spatiotemporal expression of 14 Zips and 10 ZnTs(12, 49). These transporters are located either in the plasma membrane or in the membrane of intracellular organelles where they act to modulate intracellular zinc ion by mediating its influx/efflux or release/sequestration, respectively(50). A limited number of transporters participate in the regulation of immune responses due to their tissue- or cell-specific expression. It has suggested pivotal immunoregulatory roles for Zip6 and Zip8 in T cells(6, 18). Our data illustrate that Zip1 and Zip8 are constitutively expressed by resting monocytes and macrophages. Moreover, Zip8 expression was found to be dramatically increased upon stimulation with LPS (Fig. 1C and D) as previously reported(38, 51). Knockdown of Zip8 via siRNA and real-time monitoring of zinc influx suggests Zip8 is a major mediator of zinc influx in human monocytes/macrophages (Fig. 2E and F). In addition to LPS, a variety of stimuli trigger Zip8 expression in several cell types. TNF-α and IL-1β greatly increase Zip8 expression in mouse lung epithelial and articular chondrocytes(52), respectively. TCR-stimulated T cells and IL-5-treated B cells also display enhanced Zip8 expression(18, 53), suggesting the critical role of this transporter for maintaining zinc homeostasis in cells exposed to the inflammatory milieu. Zip8 is a downstream target gene of NF-κB(38), and the re-analysis on our previous microarray data showed that expression of Zip8 is greatly upregulated in peripheral monocytes from RA patients when compared with those of healthy controls (Fig. 1A) (https://www.ebi.ac.uk/arrayexpress/experiments/E-MTAB-6187/).

RA is a prototype systemic autoimmune disease characterized by chronic inflammatory responses(54, 55). Accumulating evidence reveals that monocytes and macrophages play critical roles in the pathophysiology of RA via delivering enhanced costimulatory signaling producing pro-inflammatory cytokines(56). Moreover, it has shown that sustained hyper-inflammatory activity of glycolytic macrophages is a central parameter of RA (24, 47, 57), implying that metabolic reprogramming occurs in these cells. Our results indicate that the increase in intracellular zinc in response to Zip8 expression plays a role in inflammatory responses. Of note, peripheral and synovial monocytes of RA patients had higher mRNA expression of Zip8 and MT2A, compared with those of HCs. Considering the correlation among the mRNA expression of Zip8, MT2A, IL-1β of monocytes, and clinical parameters in patients (Fig. 7A and C), an enhanced zinc influx by monocytes from RA patients may play an important role in the regulation of their inflammatory responses. In contrast to a previous report and our findings(8), Lui *et al*. recently suggested that proinflammatory stimuli induce Zip8 expression in an NF-kB-dependent manner, resulting in zinc influx, while Zip8 negatively regulates the NF-κB pathway and proinflammatory responses through zinc-mediated suppression of IκB kinase (IKK) activity in monocytes and macrophages(38). This discrepancy in findings is likely due to the difference in cytoplasmic zinc ion levels between the two experimental systems. In our study zinc influx was caused by the increase of extracellular zinc, while in the Lui study the zinc ionophore pyrithione was utilized to promote entry of zinc ions into cells and increase cytoplasmic zinc levels. Zinc inhibits IKKα and IKKβ with an IC50 of approximately 0.5 μM. In contrast, protein tyrosine phosphatases (PTPs) are sensitive to the inhibitory action of zinc ions with IC50 values in the pM to low nM range, which is likely to be achieved *in vivo* (6). It should be noted that zinc appears to activate or inhibit several signaling pathways downstream of TLRs in human monocytes. These opposing effects are not mutually exclusive. Thus, zinc can be either pro or anti-inflammatory, depending on the concentration to which cells are exposed(37, 58).

Bioavailable zinc ions participate in regulation of many signaling pathways, and consequently, modulate immune responses(1, 40). Our data clearly illustrate that the activity of the Akt/mTOR pathway also depends on the cytoplasmic zinc concentration in human monocytes and macrophages as was previously well-depicted in the Erk1/2 pathway (Fig. 5A, B, and Suppl. Fig. 4)(8). Zinc-mediated Akt activation has been reported in several cell types including lung epithelial cells and myogenic cells in which exogenous zinc facilitates cell survival and proliferation, respectively, via largely unclear mechanisms(59, 60). Despite its essential role in immune cells, few studies have investigated the modulation of the Akt/mTOR pathway by zinc. Plum et al. recently demonstrated that zinc ions augment interleukin-2-mediated Akt phosphorylation in both a murine T cell line and thymocytes, which is due to zinc-mediated inhibition of PTEN(61). In the present study, we found that cytoplasmic zinc ions greatly enhance the phosphorylation of S6 kinase, a downstream target of mTORC1 signaling (Fig. 5B). Since mTORC1-S6K signaling serves to promote aerobic glycolysis, which is intimately connected to inflammasome activation in innate cells, we hypothesized that zinc functions as an ionic signaling molecule for the mTORC1-S6K pathway. Although S6K is a distal downstream molecule in the Akt signaling pathway, our experiments using MK-2206, a selective Akt inhibitor, strongly suggest other mechanisms underlying the zinc-dependent enhancement of S6K activity. Given that zinc is a potent inhibitor of many phosphatases, activation-dependent zinc influx may influence the activity of phosphatases directly targeting S6K.

Robust aerobic glycolysis is a hallmark of metabolic reprogramming in activated monocytes/macrophages, which is necessary to meet the demands of an immune response(24). Metabolic reprogramming of innate myeloid cells is particularly relevant for IL-1β production(24, 26, 27). mTORC1 is a key player in glycolytic reprogramming, which involves the increased translation of glycolytic enzymes or their transcriptional regulators(27). Our study clearly shows that zinc influx in activated monocytes/macrophages leads to augmented IL-1β production *via* enhanced mTORC1 activity and glycolysis (Fig. 3,4, and 5). It has been demonstrated that metabolic shift toward glycolysis largely depends on glucose uptake, which is mainly mediated by activation-induced expression of Glut1 and glucose metabolism-related enzymes. Our recent studies also suggest that the activity of mTORC1, a central regulator of glycolysis, is regulated by the intracellular level of certain amino acids, including leucine and arginine, *via* their specific sensor proteins(24) and independently of glucose utilization. In LPS-stimulated murine macrophages, mTORC1-induced hexokinase 1 (HK1)-dependent glycolysis regulates NLRP3 inflammasome activation and augments IL-1β production(27). Thus, we sought to explore the mechanism underlying the direct regulation of mTORC1-S6K activity by cytoplasmic zinc.

Protein phosphorylation is a major and reversible posttranslational modification and precisely modulated by protein kinases and phosphatases. Zinc ions are recognized as inhibitors of protein tyrosine phosphatases (PTPs) that conserve tyrosine phosphorylation and generally sustain signaling activity(6) by yet unidentified mechanisms(62). A number of studies showed that zinc binds very tightly to PTPs with an IC_50_ in the nanomolar to picomolar range. These free cytosolic zinc ion concentrations are likely achievable during signaling events in cells, thus suggesting zinc modulates PTP activity *in vivo*(23). In addition to PTPs, zinc ions also inhibit activity of several serine/threonine phosphatases including calcineurin(18). Our findings in Figure 5 suggest that the zinc-mediated enhancement of S6K activity is dependent not only on upstream signaling events, such as Akt, but also on the activity of the protein phosphatases that regulate S6K activity. PP2A and PHLPP have been identified as phosphatases that control phosphorylation and activity of S6K(43, 44, 63). Therefore, we sought to investigate whether zinc ions modulate PP2A and PHLPP activity in human monocytes/macrophages leading to changes in S6K activity.

PP2A is a highly conserved and ubiquitous serine/threonine phosphatase involved in many essential aspects of cellular function including cell cycle regulation, cell growth control, and signal transduction pathways(64, 65). The PP2A complex consists of three subunits, a structural regulatory subunit (A), a regulatory subunit (B), and a catalytic subunit (C). PP2A was initially identified as an important tumor suppressor protein due to its relationship with cell cycle and cell growth(66). However, accumulating evidence suggests involvement of PP2A in the control of inflammation. Treg cell-specific loss of PP2A leads to altered metabolic and cytokine profiles and impaired capability to suppress immune responses. Further, Th17 differentiation is dependent on PP2A, through its involvement in regulation of the canonical TGFβ-SMADs-RORγt signaling process(42, 67). As seen in Figure 6B and C, LB-100, a small molecule inhibitor of the PP2A-C subunit, enhances phosphorylation of S6K and production of IL-1β, demonstrating that PP2A functions as a phosphatase regulating S6K activity. Early studies showed that THP-1, a human monocytic T cell line, constitutively expresses all three subunits of PP2A. In line with this, PP2A levels were unaffected by LPS stimulation in our study (Fig. 6A) and PP2A regulated JNK activity via a direct physical interaction in the context of an inflammatory stimulus. More recently, myeloid-specific deletion of PP2A-Cα in murine BMDMs was found to intensify MyD88- and TRIF-dependent inflammation following LPS challenge, suggesting an important regulatory role of PP2A in many aspects of inflammation and survival in activated myeloid cells (68). Of importance, studies have suggested that PP2A-B, the regulatory subunit, interacts with and dephosphorylates S6K, whose activity was found to be dramatically influenced by intracellular zinc concentration in the current study (Fig. 5). PP2A was found to physically interact with overexpressed S6K in 293 T cells and with endogenous S6K in primary human macrophages, similar to what was reported in *Drosophila*(44). Considering that phosphorylation of the PP2A-C subunit at Tyr307 is a negative indicator of PP2A activity (46, 69), our data suggest that the zinc-mediated enhancement of S6K activity is attributable to PP2A in monocytes/macrophages (Fig. 6F).

Our findings have clinical importance for the unchecked or prolonged mTORC1-induced glycolysis in several chronic inflammatory disorders(24). Activation-induced upregulation of Zip8 expression and enhanced zinc influx could explain metabolic shift to glycolysis seen in monocytes from patients with RA. Considering the correlation between MT2A and IL-1β levels in circulating monocytes of RA patients (Fig. 7), Zip8-mediated zinc influx by monocytes and macrophages could be an important role for pathogenesis of chronic inflammatory disorders. Thus, Zip8 may be a potential therapeutic target for a variety of inflammatory disorders

In conclusion, we provide evidence that cytoplasmic bioavailable zinc is important for modulation of IL-1β production in human monocytes and macrophages. Upon stimulation, cytoplasmic levels of bioavailable zinc in these cells is largely influenced by extracellular zinc concentrations, in part *via* Zip8-mediated influx. The phosphorylation of S6 kinase is enhanced and maintained under increased zinc concentrations via zinc-mediated inhibition of PP2A, an S6K phosphatase, and as a result, IL-1β production is increased by activated mTORC1-induced glycolysis. In RA patients, the expression of Zip8 and MT2A by peripheral and synovial monocytes was significantly increased and their Zip8 levels positively correlated with RA clinical parameters, suggesting that Zip8-mediated zinc influx is related to inflammatory conditions. Our data provide new insight into how bioavailable zinc modulates cytokine production in human monocytes/macrophages *via* metabolic reprogramming which is an essential process for inflammatory responses.

## Materials and Methods

### Cell preparation

The study protocols were approved by the institutional review board of Seoul National University Hospital, Chungnam National University Hospital, and Severance Hospital, Yonsei University Health System. Peripheral blood of RA patients and healthy donors was drawn after obtaining written, informed consent. The methods were performed in accordance with the approved guidelines. Peripheral blood mononuclear cells (PBMCs) were isolated from blood by density gradient centrifugation (Bicoll separating solution; BIOCHROM Inc., Cambridge, UK). Monocytes were positively separated from PBMCs with anti-CD14 magnetic microbeads (Miltenyi Biotec Inc., Auburn, CA, USA).

### Cell culture

Purified monocytes were cultured in serum free RPMI 1640 medium supplemented with 1% penicillin/streptomycin and 1% L-glutamine. Human monocyte-derived macrophages (HMDMs) were differentiated from purified CD14^+^ monocytes in the presence of recombinant human M-CSF (50 ng/ml; PeproTech, Cranbury, NJ, USA) for 6 days in RPMI 1640 medium supplemented with 10% FBS, 1% penicillin/streptomycin and 1% L-glutamine.

### Antibodies and reagents

LPS, TPEN (N,N,N’,N’-tetrakis-(2-pyridyl-methyl)ethylenediamine), ATP, Glucose, oligomycin, 2-DG (2-deoxy-D-glucose), and Okadaic acid were obtained from MilliporeSigma (Burlington, MA, USA). LB100, PP2A-specific small molecule inhibitor, was purchased from Selleckchem (Houston, TX, USA). Anti-IL-1β, anti-caspase 1, anti-phospho p70 S6 Kinase (Thr389), anti-p70 S6K, anti-phospho mTOR (Ser2448), anti-mTOR, anti-phospho Akt (Ser308 and Ser473), anti-Akt, anti-phospho 4EBP1 (Ser65), anti-4EBP1, anti-phospho NF-κB p65 (Ser536), anti-NF-κB p65, anti-phospho ERK (Thr202/Tyr204), anti-ERK, anti-phospho p38 (Thr180/Tyr182), anti-p38, and anti-PP2A-B subunit antibodies (Abs) were purchased from Cell Signaling Technology (Danvers, MA, USA). Anti-PP2A-C and anti-β-actin Ab was obtained from MilliporeSigma (Burlington, MA, USA). Phospho-PP2A-C (Tyr307) Abs were purchased from Santacruz (Dallas, Texas, USA), respectively.

### Enzyme-Linked Immunosorbent Assay (ELISA)

The amount of IL-1β, TNF-α and IL-6 in culture supernatant was quantified by commercial ELISA kits (Thermo Fisher Scientific, Waltham, MA, USA). The measurement of OD (Optical density) was performed using the infinite 200 pro multimode microplate reader (Tecan Group Ltd., Seestrasse, Switzerland).

### Immunoblot Analysis

Monocytes and macrophages were lysed in RIPA lysis buffer (150 mM NaCl, 10 mM Na_2_HPO_4_, pH 7.2, 1% Nonidet P-40, and 0.5% deoxycholate) containing PMSF (phenylmethylsulfonyl fluoride) (MilliporeSigma), EDTA, and protease and phosphatase inhibitor cocktail (Thermo Fisher scientific). Proteins from supernatants were precipitated using methanol/chloroform. Cell lysates were separated on 8-12% SDS-PAGE gel and transferred onto a PVDF membrane (Bio-Rad, Hercules, CA, USA). The membrane was incubated overnight with the respective primary antibodies at 4 °C, and then incubated with peroxidase-conjugated secondary Abs for 1 h (Cell signaling) for 1 h at room temperature. The membranes were developed by ECL system.

### Immunoprecipitation

Cell lysates were prepared using modified RIPA buffer (50 mM Tris-HCl, pH 7.4, 150 mM NaCl, 1% Nonidet P-40, and 0.25% deoxycholate) containing PMSF, EDTA, and protease and phosphatase inhibitor cocktail. Cell lysates (500 μg) were incubated with target antibodies at 4°C for overnight and immunoprecipitated with protein A/G plus agarose beads (Santacruz) for 2 hr at 4°C.

### RT-PCR

Total RNA was extracted from freshly isolated or cultured cells using TRIzol reagents (life technologies, Grand Island, NY, USA), and cDNA was synthesized by GoScript reverse transcription system (Promega, Madison, WI, USA). Real-time quantitative RT-PCR was performed in duplicates on a CFX Automation System (Bio-rad). The levels of gene expression were normalized to the expression of β-actin. The comparative Ct method (*ΔΔ*Ct) was used for the quantification of gene expression.

### Intracellular zinc measurements with fluorescent probes

Cells were incubated in loading buffer [HBBS (-), 1 mM Ca^2+^, 1 mM Mg^2+^, 0.5% BSA] for 30 min, either with 1 μM FluoZin-3-AM (Thermo Fisher scientific) at 37°C. After cells were washed twice with washing buffer [PBS + 10% Bovine Serum + 1% Penicillin/Streptomycin], the fluorescence was recorded on infinite 200 pro multimode microplate reader using excitation and emission wavelengths of 485 and 535 nm. The concentration of intracellular zinc was calculated from the mean fluorescence with the formula [Zn] *= Kd* ⅹ [(*F*-*F*_min_)]/(*F*_max_-*F*)], using 50 μM TPEN to determine minimal fluorescence and 100 μM ZnCl_2_/50 μM pyrithione to determine maximal fluorescence, respectively (70).

### Metabolic Analysis

Human primary monocytes and macrophages were pre-treated with TPEN 1.5 µM or ZnCl_2_ (0 and 45 µM) and stimulated with 10 or 100 ng/ml LPS for 24 h in RPMI 1640 medium supplemented with 10% fetal bovine serum, 1% penicillin/streptomycin, and 1% l-glutamine. To measure cellular respiration activity of the cells, monocytes were seeded as a monolayer onto XFe24 cell culture plates (Seahorse Bioscience, MA, USA). The culture media was replaced with XF assay media supplemented with l-glutamine (250 μg/ml) and incubated for 1 h in non-CO2 incubator. Glucose (10 mM), oligomycin (2 μM), and 2-DG (50 mM) were sequentially treated into the cells during real-time measurements of extracellular acidification rate (ECAR) and Oxygen consumption rate (OCR) using XFe24 analyzer. Glycolysis parameters were calculated using XF glycolysis stress test report generator program that was provided from manufacturer (Seahorse Bioscience). Glycolysis and glycolysis capacity were calculated by subtracting ECAR after glucose treatment from ECAR before oligomycin and subtracting ECAR after oligomycin from ECAR before 2-DG treatment, respectively.

### Lactate production assay

The lactate production is measured using colorimetric assay (BioVision Technologies, Milpitas, CA, USA), according to the manufacturer’s instructions. Absorbance was measured using infinite 200 pro multimode microplate reader at 570 nm.

### Zinc assay

Zinc levels were estimated using a commercially available zinc assay kit (MilliporeSigma), according to the manufacturer’s instructions. Absorbance was measured using infinite 200 pro multimode microplate reader at 560 nm.

### Phosphatase assay

PP2A activity in cell lysates was measured using the Phosphatase Assay Kit (Promega). Endogenous free phosphate was removed using the columns, and then the extracts were normalized for protein content. Lysates (5 μg) were incubated with diverse ZnCl_2_ or PP2A inhibitors for 30 min at 30 °C using a Thermo-mixer (Eppendorf, Hamburg, Germany). The protein samples were incubated for 30 min at 33 °C with a chemically synthesized phosphopeptide [RRA(pT)VA], as a substrate for PP2A, PP2B, and PP2C, not for PP1 in optimized buffer for PP2A activity while cation-dependent PP2B and PP2C were inhibited. Released phosphate from substrate was detected by adding an equal volume of the Molybdate Dye/Additive mixture for 15 min at 630 nm. PP2A activity was calculated by the release of phosphate per μg of protein and per minute (pmol/μg/min), according to the manufacturer’s instructions.

### Statistics

A paired t-test, unpaired t-test, or Pearson correlation analysis was done to analyze data using Prism 7 software (GraphPad Software Inc., La Jolla, CA, USA) as indicated in the figure legends. *p* -values of less than 0.05 were considered statistically significant.

## SUPPLEMENTARY MATERIALS

**Supplementary Figure 1.** Enhanced expression of zinc transporter Zip8 in monocytes derived in synovial fluid (SF) of RA patients compared to those in their peripheral blood.

**Supplementary Figure 2.** Knockdown of Zip8 in human monocyte-derived macrophages (HMDMs).

**Supplementary Figure 3.** Effect of zinc on production of proinflammatory cytokines.

**Supplementary Figure 4.** Increased intracellular zinc leads to upregulated phosphorylation of Akt/mTORC1 signaling pathway molecules.

**Supplementary Figure 5.** Role of PP2A in zinc-mediated regulation of S6K activity in human monocytes.

**Supplementary Figure 6.** PHLPP does not play a role in zinc-mediated regulation of S6K activity in human monocytes and macrophages.

**Supplementary Figure 7.** Clinical relevance of zinc-mediated metabolic reprograming in monocytes of RA patients.

## ACKNOWLEDGMENTS

The authors thank Jiyeon Jang (Seoul National University College of Medicine) for assisting in the recruitment of human subjects and thank Core Lab, Clinical Trials Center, Seoul National University Hospital for drawing blood.

## AUTHOR CONTRIBUTIONS

B.K: participated in the design of the study, performed most of the experiments, data collection and analysis, and drafted manuscript. H.Y.K. and B.R.Y.: participated in the design of the study, performed the experiments, data collection and analysis. J.Y., K.-S.Y., H.C.K., J.K.P. and S.W.K.: participated in its design and performed data analysis. W-W.L.: conceived of the study, participated in its design and coordination, performed data analysis and writing of manuscript, and has full access to all the data in this study and financial support. All authors have read and approved the final manuscript.

