## Supplementary figure 1-7 for "Cytoplasmic zinc regulates IL-1β production by monocytes/macrophages via mTORC1-induced glycolysis in rheumatoid arthritis (RA)"

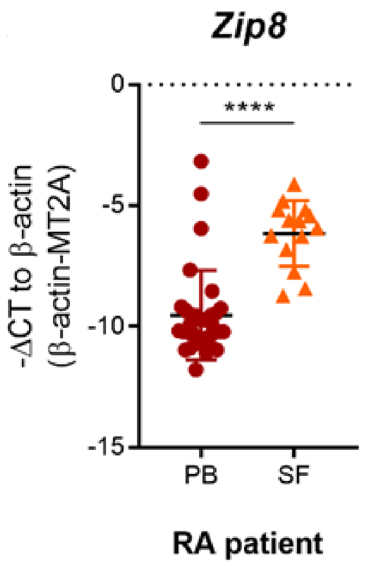


**Supplementary Figure 1. Enhanced expression of zinc transporter Zip8 in monocytes derived in synovial fluid (SF) of RA patients compared to those in their peripheral blood.** mRNA level of Zip8 was quantified by real-time RT-PCR in peripheral monocytes of peripheral ($n$=32) and synovial monocytes ($n$=13) of RA patients. Bar graphs show the mean ± SD. **** = $p < 0.0001$ by unpaired $t$-test.

#### Supplementary Figure 2

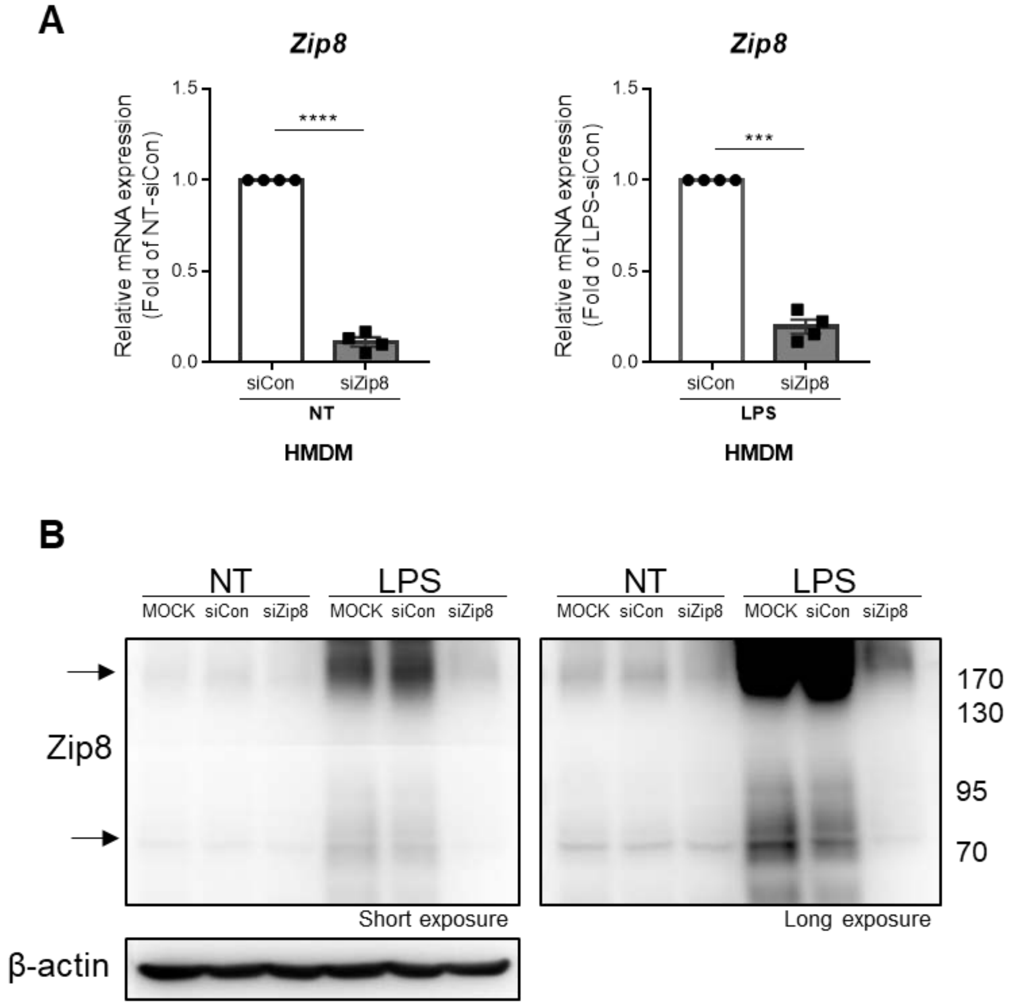


**Supplementary Figure 2. Knockdown of Zip8 in human monocyte-derived macrophages (HMDMs).** Macrophages were transfected with Zip8-targeted or control siRNA (20 pM of both siRNAs). At 24 h post-transfection, siRNA-transfected cells were stimulated with LPS for 24 h. (**A-B**) The efficiency of Zip8-specific knockdown was evaluated by its mRNA (A) and protein (arrows in B) expression. Data is representative of three independent experiments with three different donors. Bar graphs show the mean ± SEM. *** = $p < 0.001$ and **** = $p < 0.0001$ by paired *t*-test.

### Supplementary Figure 3

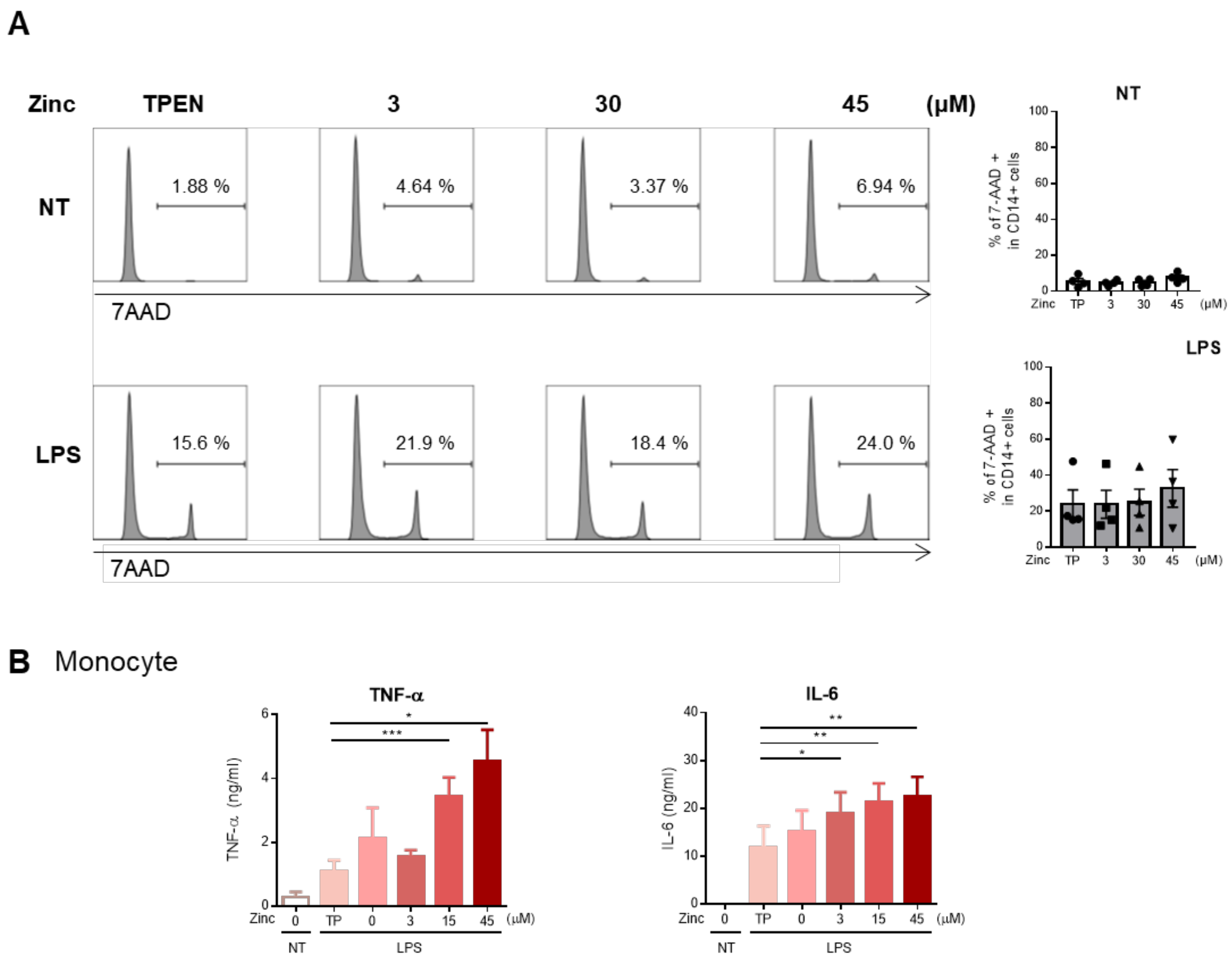


**Supplementary Figure 3. Effect of zinc on production of proinflammatory cytokines.** Human primary monocytes were incubated with zinc chelator TPEN (150 nM) or the indicated concentration of $ZnCl_2$ for 2 hr, followed by stimulation with LPS (10 ng/ml) for 24 hr. **(A)** The rate of cell death was assessed by 7-AAD staining in the presence of TPEN or various concentrations of zinc. **(B)** The amount of TNF-α and IL-6 in the supernatants was quantified by ELISA (n=7). The amount of IL-1β in the supernatants was quantified. Bar graphs show the mean ± SEM. * = $p < 0.05$, ** = $p < 0.01$ and *** = $p < 0.001$ by two tailed paired *t*-test.

### Supplementary Figure 4

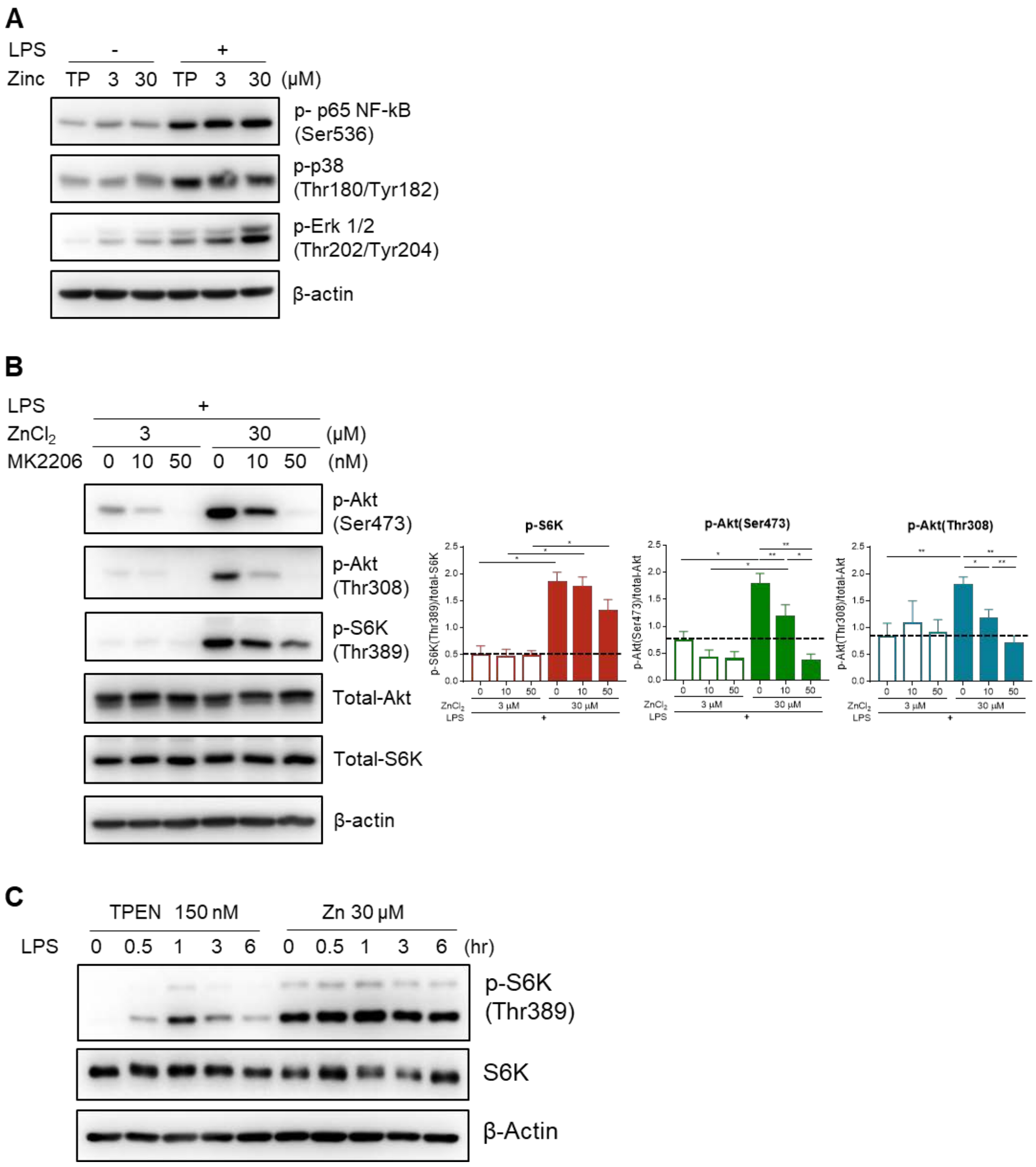


**Supplementary Figure 4. Increased intracellular zinc leads to upregulated phosphorylation of Akt/mTORC1 signaling pathway molecules. (A)** Human primary monocytes were treated with TPEN (150 nM) or $ZnCl_2$ for 2 hr and stimulated with or without LPS (10 ng/ml) for 15 min. Cell lysate were prepared and immunoblotted for the indicated signaling molecules (n=3). **(B)** Human primary monocytes were pre-treated with the indicated concentrations of MK2206 and incubated with $ZnCl_2$ (3 or 30 μM) for 30 min, followed by stimulation with LPS for 15 min. Cell lysate were prepared and immunoblotted for the indicated signaling molecules. The inhibitory effect of MK2206 on the phosphorylation of Akt and S6K in monocytes treated with 3 or 30 μM $ZnCl_2$ was plotted with band intensities. The phosphorylation level was normalized to the total protein for each signaling protein. **(C)** Macrophages were pre-incubated with TPEN or $ZnCl_2$ for 2 hr, followed by stimulation with LPS for the indicated time. β-actin was used as a normalization control. Bar graphs show the mean ± SEM. * = $p < 0.05$ and ** = $p < 0.01$ by two-tailed paired *t*-test.

#### Supplementary Figure 5

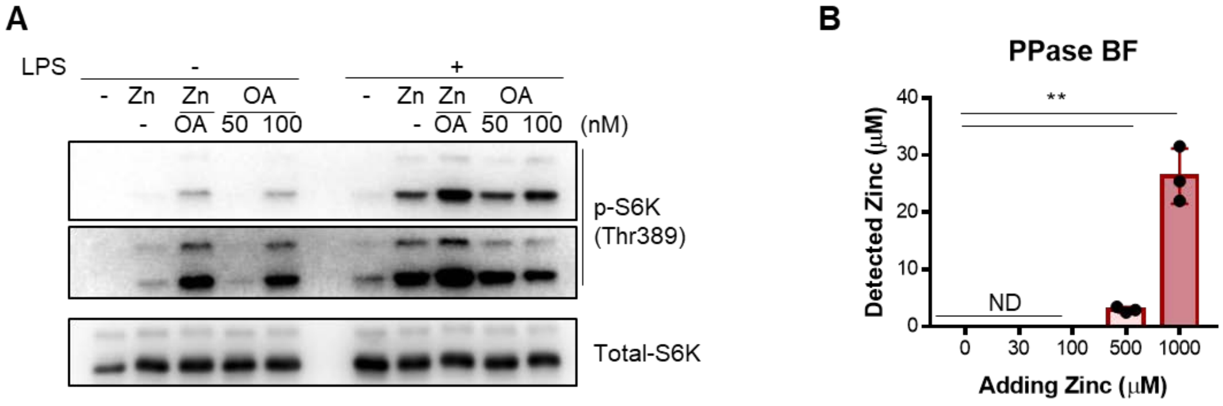


**Supplementary Figure 5. Role of PP2A in zinc-mediated regulation of S6K activity in human monocytes. (A)** Monocytes were pretreated for 2 hr with Okadaic acid (OA: general PP2A and PP1 inhibitor) with or without $ZnCl_2$ (30 μM). Cells were stimulated for 15 min with or without LPS and their lysates were immunoblotted for total S6K and p-S6K. **(B)** Zinc ion concentration was measured in the buffer for the phosphatase protein 2A assay treated with the indicated concentration of $ZnCl_2$. Bar graphs show the mean ± SEM. ** = $p < 0.01$ by two tailed paired *t*-test.

#### Supplementary Figure 6

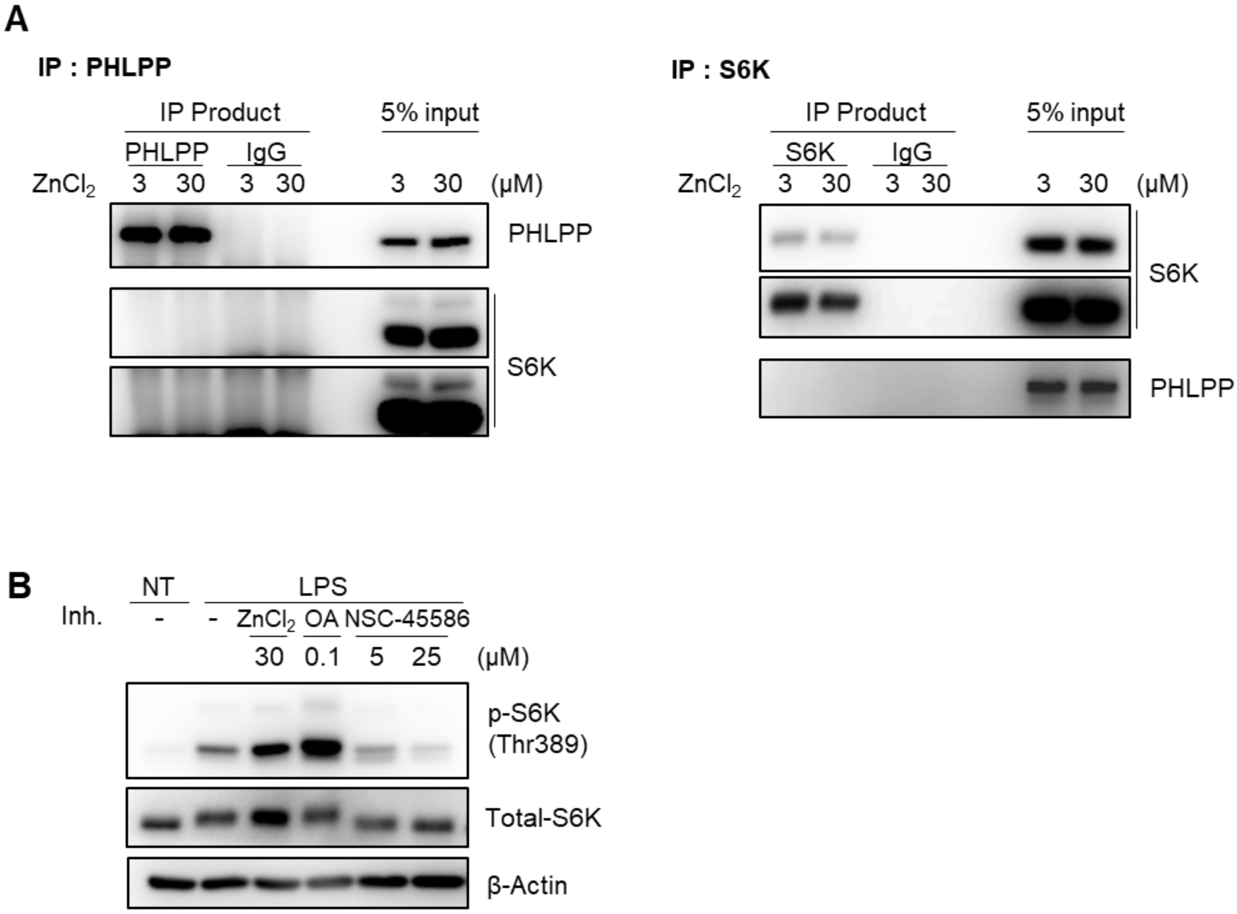


**Supplementary Figure 6. PHLPP does not play a role in zinc-mediated regulation of S6K activity in human monocytes and macrophages. (A)** THP-1 derived macrophages were treated with $ZnCl_2$ at 3 or 30 μM. Cell lysates were immunoprecipitated with Abs to PHLPP or S6K and immunoblotted with the indicated Abs **(B)** Monocytes were pretreated for 2 hr with $ZnCl_2$ at 30 μM, Okadaic acid (OA: general PP2A and PP1 inhibitor) or NSC-45586 (PHLPP inhibitor). Cells were stimulated for 15 min with or without LPS and their lysates were immunoblotted for total S6K and p-S6K.

#### Supplementary Figure 7

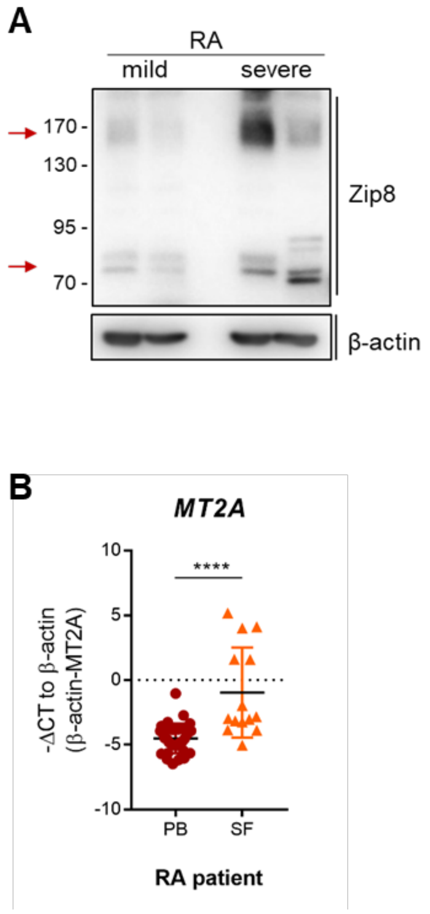


**Supplementary Figure 7. Clinical relevance of zinc-mediated metabolic reprograming in monocytes of RA patients. (A)** Cell lysates of peripheral monocytes of RA patients were prepared for immunoblotting for Zip8 proteins. The values of clinical parameters in each patient of two groups [ESR DAS28: 2.83 and 0.91 of mild vs. 6.05 and 7.47 of severe group; CRP DAS28: 1.36 and 1.45 of mild vs. 4.56 and 5.97 of severe group]. **(B)** mRNA level of MT2A was quantified by real-time RT-PCR in peripheral monocytes of peripheral (*n*=32) and synovial monocytes (*n*=13) of RA patients. Bar graphs show the mean ± SD. **** = $p < 0.0001$ by unpaired *t*-test.
